# Unsupervised learning of progress coordinates during weighted ensemble simulations: Application to millisecond protein folding

**DOI:** 10.1101/2024.08.28.610178

**Authors:** Jeremy M. G. Leung, Nicolas C. Frazee, Alexander Brace, Anthony T. Bogetti, Arvind Ramanathan, Lillian T. Chong

## Abstract

A major challenge for many rare-event sampling strategies is the identification of progress coordinates that capture the slowest relevant motions. Machine-learning methods that can identify progress coordinates in an unsupervised manner have therefore been of great interest to the simulation community. Here, we developed a general method for identifying progress coordinates “on-the-fly” during weighted ensemble (WE) rare-event sampling via deep learning (DL) of outliers among sampled conformations. Our method identifies outliers in a latent space model of the system’s sampled conformations that is periodically trained using a convolutional variational autoencoder. As a proof of principle, we applied our DL-enhanced WE method to simulate a millisecond protein folding process. To enable rapid tests, our simulations propagated discrete-state synthetic molecular dynamics trajectories using a generative, fine-grained Markov state model. Results revealed that our on-the-fly DL of outliers enhanced the efficiency of WE by >3-fold in estimating the folding rate constant. Our efforts are a significant step forward in the unsupervised learning of slow coordinates during rare event sampling.

## 1 Introduction

Rare-event sampling methods have been increasingly used to simulate long-timescale biological processes at the atomic level.^1–3^ For many of these methods, a major challenge that remains is the identification of a progress coordinate (also known as reaction coordinate or collective variables) that captures the relevant slow motions. Given that the intrinsic dimensionality of a molecular dynamics (MD) simulation with N atoms is 3N-6 (in Cartesian coordinates), even relatively small systems can be challenging to analyze using approaches that focus on motions along only a few dimensions. Strategies for identifying progress coordinates include the use of fast, approximate trajectories,^4^ identification of coordinates that correlate with the committor (or commitment probability),^5,6^ and automated artificial intelligence (AI) techniques such as machine and deep learning.^7–12^

AI techniques can identify progress coordinates by detecting distinct conformational states in an unsupervised manner based solely on the atomic coordinates of structures sampled by an MD simulation. This detection is commonly facilitated by projecting the high-dimensional data from MD simulations onto low-dimensional manifolds containing a compressed representation of data. As demonstrated by a recent study, deep learning (DL) techniques can identify effective progress coordinates for simulating the folding of small proteins via analysis of extensive MD simulations and the use of such progress coordinates with adaptive sampling has accelerated the sampling of folding events by >100x relative to conventional MD (cMD) simulations. ^7^

Here, we have developed a DL method to learn progress coordinates “on-the-fly” during weighted ensemble (WE)^13–15^ rare-event sampling. ^16–18^ WE is a path sampling strategy that has enabled atomistic simulations of complex processes such as protein folding, ^19^ protein-ligand (un)binding,^20^ and large-scale conformational transitions in proteins.^21^ The DL method involves applying a convolutional variational autoencoder (CVAE) to compress high-dimensional WE simulation data down to lower-dimensional representations in latent space and then replicating outlier trajectories during a resampling procedure (Figure 1). CVAE models are particularly effective in anomaly detection through capturing spatial relationships between the pixels of an image.^22^

**Figure 1:**
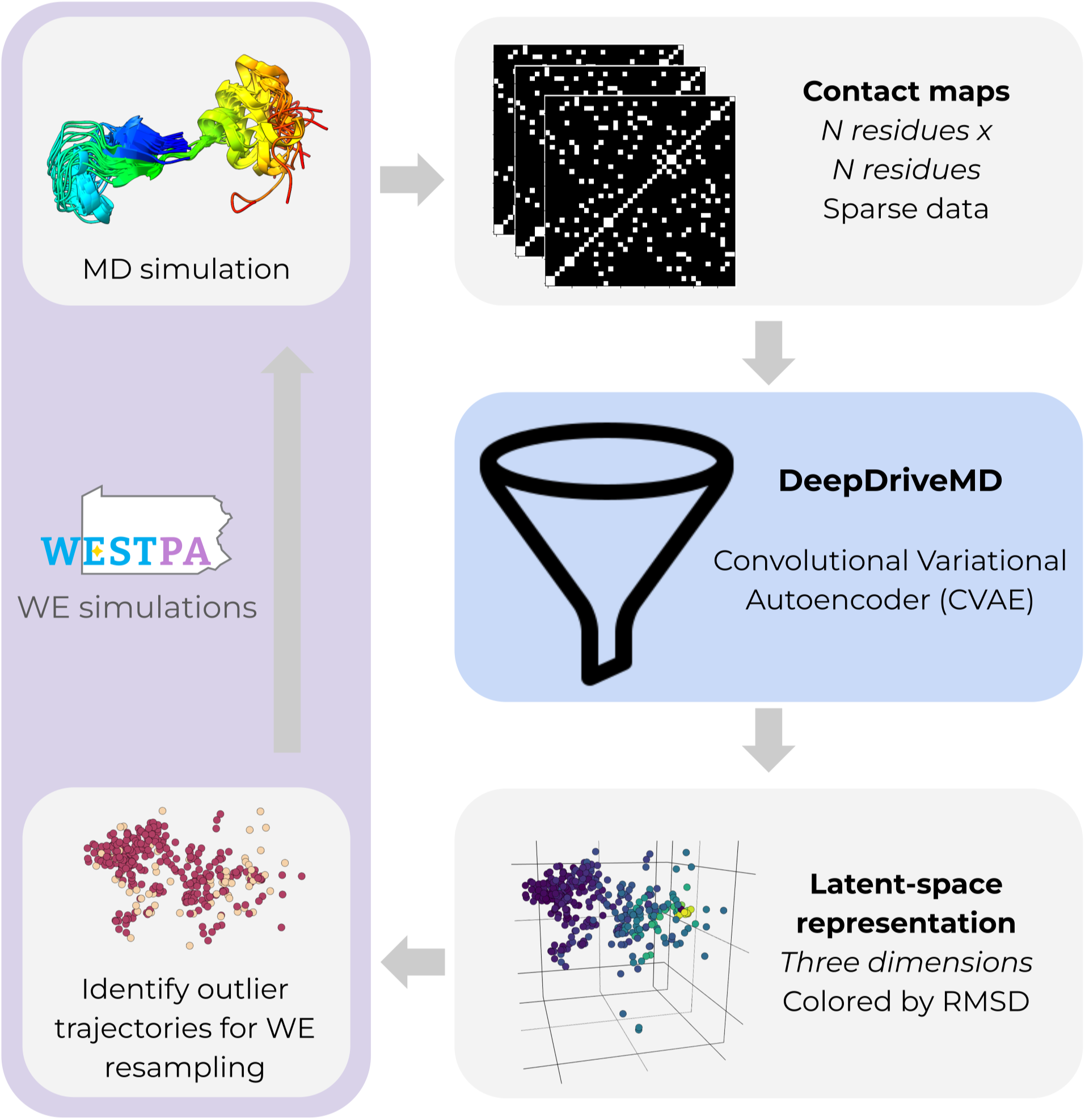
Workflow for DL-enhanced WE simulations. On-the-fly DL of progress coordinates during a WE simulation involves applying a convolutional variational autoencoder (CVAE) to compress the high-dimensional simulation data down to a three-dimensional latent space model. The high-dimensional data is in the form of pairwise residue contact matrices for selected conformations from the WE simulation. A WE resampling procedure is periodically applied by replicating outlier trajectories to enrich for sampling of rare, barriercrossing transitions (e.g., protein folding). WE simulations are run using the WESTPA software^17,18^ and the CVAE model^22^ is created using the DeepDriveMD software.^13–15^

As a proof of principle, we applied our DL-enhanced WE strategy to simulate the millisecond folding process of the N-terminal domain of L9 (NTL9) protein. Our simulations employed discrete-state synthetic molecular dynamics (synMD) trajectories,^23^ which are ideal for methods testing due to their greatly reduced computational cost, atomistic structures, and analytical “ground-truth” solution for steady-state observables (i.e., rate constants). We determine the features of the simulation data that are needed to build an effective latent space model of the system and train the latent space model “on-the-fly” to learn an effective progress coordinate for the molecular process of interest.

## 2 Methods

### Overview of WE path sampling

WE path sampling enhances the sampling of pathways and rates for rare events such as protein folding and binding by running a large number *M* of weighted trajectories in parallel and iteratively applying a resampling procedure at fixed time intervals τ.^16,24^ At each WE iteration, the resampling procedure involves replicating trajectories that have occupied less-visited regions of configurational space and occasionally terminating trajectories that have occupied more frequently visited regions. Such regions are typically defined as bins or clusters along a progress coordinate. Trajectory weights are tracked rigorously such that the weights sum to a total probability of one thereby ensuring that no bias is introduced into the dynamics. To maintain a non-equilibrium steady state, trajectories that reach the target state (e.g., folded state for protein folding) are “recycled”, initiating a new trajectory from the initial state (e.g., unfolded state) with the same statistical weight.

### DL-enhanced WE simulations of protein folding

To further enhance the efficiency of WE simulations in sampling rare events, we have developed a method that employs DL to learn progress coordinates on-the-fly during a WE simulation. All WE simulations were run using the WESTPA 2.0 software (https://github.com/westpa/westpa),^17^ in conjunction with synMD trajectories^23^ DL analysis was carried out using the mdlearn Python library associated with the DeepDriveMD software (https://github.com/ramanathanlab/ mdlearn).^13–15,25^ The mdlearn library includes linear, non-linear, and hybrid machine learning tools for learning latent space representations (embedding models) of MD simulation data to characterize biologically relevant conformational transitions. ^26–30^ While the DeepDriveMD software orchestrates adaptive sampling using various MD engines, the mdlearn library provides support for ML/AI methods within the DeepDriveMD software.^14,15^

In our workflow for DL-enhanced WE simulations (Figure 1), the DeepDriveMD software compressed high-dimensional pairwise residue contact maps down to three-dimensional, latent space representations using a convolutional variational autoencoder (CVAE).^22^ In the contact maps, a pair of residues was considered to be in contact if the minimum distance between their C_α_ atoms was within 8 Å. The DL-enhanced WE resampling procedure was focused on the replication of trajectories from selected “outlier” conformations, which were identified by applying the Local Outlier Factor (LOF) anomaly detection method^31^ to CVAE latent space representations of trajectory data and monitoring a single structural feature of the protein system in real space, i.e. C_α_ RMSD from the folded structure. The LOF method, as implemented in scikit-learn,^32^ is an unsupervised learning algorithm used to identify outliers in a dataset based on high deviations in local density (LOF score). Here, we sorted trajectories at the current WE iteration by the LOF score, which was based on each conformation’s local neighborhood containing 20 neighboring points calculated among up to 1000 other conformations that were randomly selected from all past WE iterations. We then replicated trajectories with high LOF scores (outliers from undersampled regions) and terminated trajectories with low LOF scores (inliers from oversampled regions). For calculation of the LOF scores, distances between any two conformations A and B were calculated using a normalized Euclidean distance metric between their latent space coordinates 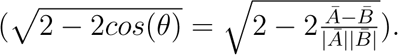. The C RMSD from the folded structure was calculated in real space using regular Euclidean distances 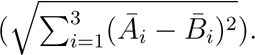.

At each WE iteration, the DL-enhanced WE resampling procedure was applied in two stages (Figure 2). In the first stage, we identified outliers among the M total trajectories at the current WE iteration by (i) sorting the trajectories by LOF score, (ii) designating the top 12 of the trajectories as “outliers” (high LOF scores) and the bottom 12 as “inliers”, and (iii) ranking each list of trajectory candidates by the C_α_ RMSD from the folded structure. To avoid generating trajectories with extremely low weights, trajectories with statistical weights beyond a minimum threshold of 10^-40^ were removed from the list of outliers. Likewise, to avoid a single trajectory with a majority of the total probability, a trajectory with a statistical weight beyond a maximum threshold of 0.1 was removed from the list of inliers. In the second stage, we applied the WE resampling procedure, replicating and terminating trajectories to maintain a fixed total number of *M* =72 trajectories. Candidates for replication were the six outliers with the lowest C_α_ RMSD values and candidates for termination were the 12 inliers with the largest C_α_ RMSD values. The combination of replication and termination instances was randomly selected. For example, to yield four replicas, we may select two trajectories and replicate each trajectory once or select one trajectory and replicate that trajectory three times. The maximum combined number of replication and termination instances was set to 12.

**Figure 2:**
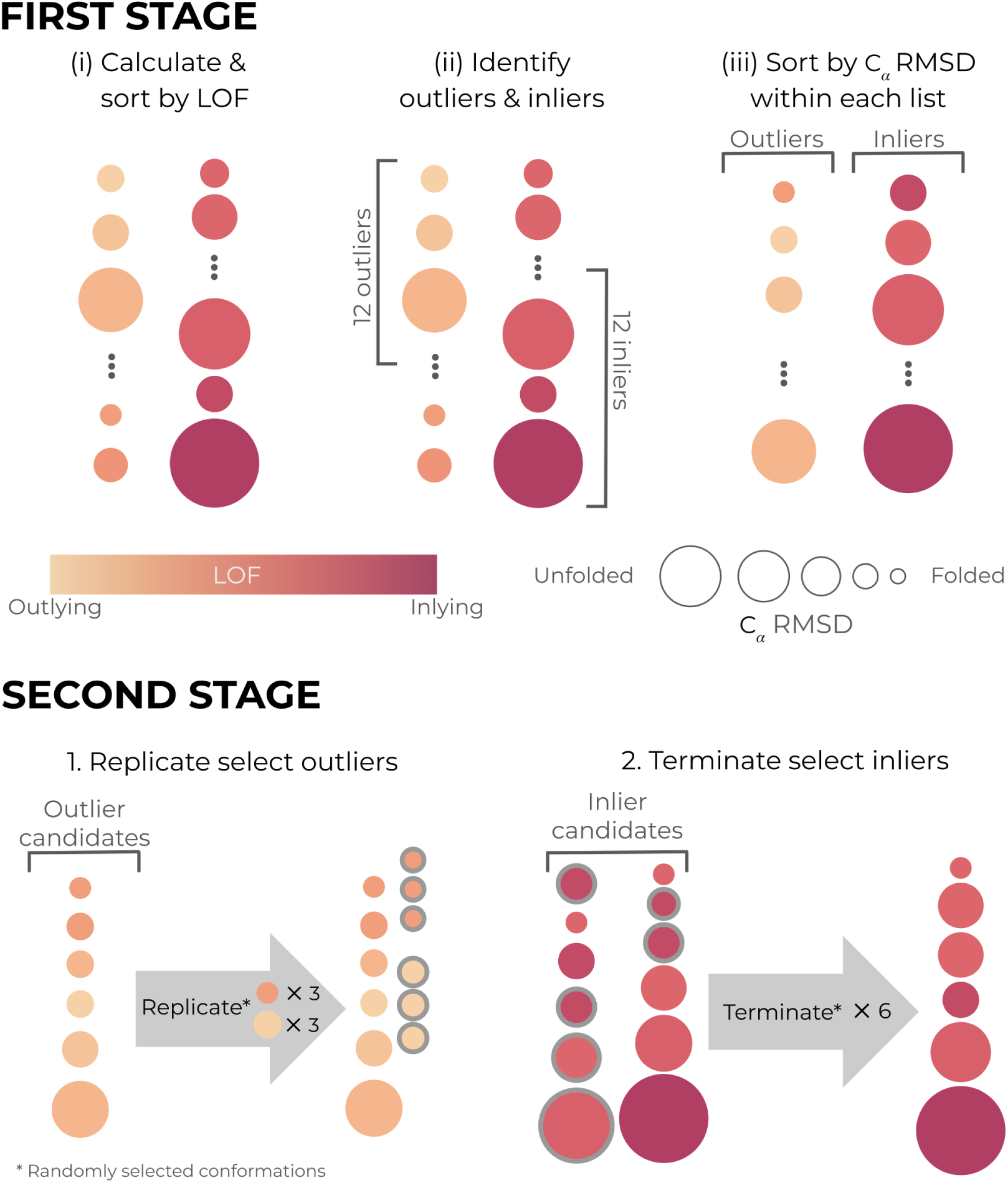
Illustration of the DL-enhanced WE resampling procedure. The resampling procedure was applied in two stages. In the first stage, trajectories were sorted by their LOF score, designating the top 12 trajectories as “outliers” and bottom 12 trajectories as “inliers”. In the second stage, a fixed number of *M* =72 total trajectories was maintained through a random combination of replicating the six lowest-RMSD trajectories and terminating the 12 highest-RMSD trajectories. The maximum combined number of replication and termination instances was set to 12. As with regular WE, trajectory weights were rigorously tracked throughout the simulation.

### Propagation of synMD trajectories

To enable rapid testing of each WE protocol with an atomistic system, we used the synMD approach^33^ to propagate discrete-state trajectories in a WE simulation. This approach involves propagating discrete-state Markov chain trajectories with a fixed time step among the “microbins” of a generative, fine-grained Markov state model (MSM) based on bin-to-bin transition probabilities. Here, our MSM was based on a set of cMD simulations of the NTL9 protein folding process (2.5 µs of total simulation time) with a lag time of 10 ps. These simulations employed the Amber ff14SB force field^34^ with generalized Born implicit solvent (Hawkins, Cramer, Truhlar model; ^35,36^ igb=1),^37^ and were performed in the NVT ensemble at 300K using a weak Langevin thermostat (collision frequency of 5 ps^-1^).

The MSM was previously constructed by Russo & Zuckerman^38^ by first generating pairwise heavy-atom distance matrices of the simulation data, excluding nearest neighbors, and then applying the variational approach for Markov processes (VAMP)^39^ to reduce the dimensions of these matrices into 356 components covering 85% of the variance. Microbins of the MSM were generated by applying a stratified k-means clustering^33^ of the simulation data in which trajectories were clustered within “strata” bins defined along their C_α_ RMSD to a reference folded structure. Any microbins that did not involve any direct or indirect microbin-to-microbin transitions to the unfolded or folded states were removed and their corresponding structures were reassigned to nearby surviving microbins. The resulting MSM consisted of 3512 microbins computed using a 10-ps lag time, which was reduced from 13,250 initial clusters (250 clusters per stratum). Stratified bin boundaries were positioned at 0.1 Å increments along [1.1, 4.5], 0.2 Å increments along [4.6, 6.4], and 0.3 Å increments along [6.6, 9.6]. The unfolded state was defined as having *≥* 9.6 Å C_α_ RMSD from a reference folded crystal structure (PDB 2HBB).^40^ The folded state was defined as having < 1 Å C_α_ RMSD from the same reference structure.

For our WE simulations of NTL9 protein folding, synMD trajectories were propagated among the 3512 microbins of the MSM mentioned above using a resampling time interval τ of 10 ps. At each τ, the microbin that was visited by a trajectory was back-mapped to a representative structure of that microbin (k-means cluster) to generate discrete trajectories of the NTL9 folding process. To maintain non-equilibrium steady state conditions, trajectories reaching the target folded state were “recycled” by initiating a new trajectory from randomly selected conformation of the initial unfolded-state ensemble with the same statistical weight. The unfolded state consisted of 22 representative conformations and the folded state consisted of one structure.

### Training of convolutional variational autoencoder models

The variational autoencoder (VAE) is a deep neural network architecture that can be used for unsupervised learning of a continuous latent-variable model that captures salient features of a dataset.

A VAE consists of an encoder *q_ϕ_*(**z***|***x**^(^*^i^*^)^) that compresses input data **x**^(^*^i^*^)^ into a small latent code **z** and a decoder *p_θ_*(**x**^(^**^i^**^)^*|***z**) that reconstructs the code to its original form.^41^ VAEs are trained on a joint optimization objective function that attempts to minimize the reconstruction error of the input data and maximize the correspondence to a selected prior distribution *p_θ_*(**z**) (e.g., Gaussian) by computing the Kullback–Leibler (KL) divergence which acts as a regularizer, via the loss function:

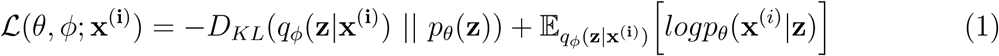

In this work, we employed a convolutional variational autoencoder (CVAE) containing four convolution layers each with 16 filters and a kernel size of 3 in the encoder, which reduced the 40 x 40 input contact matrix into a three-dimensional information bottleneck forming a latent space representation of the trajectory data. The decoder module mirrored the encoder using a series of transposed convolutions. A rectified linear-unit (ReLU) activation function was used between each interior layer, transforming the final layer output via a sigmoid activation function. As the contact map elements are binary, we computed the reconstruction loss by taking the binary cross-entropy loss of the predicted sigmoidal outputs. To regularize the model, we used a standard normal-distribution *N* (0, 1) prior for which a closed-form Kullback-Liebler (KL) divergence was derived. ^41^ The model was trained using the RMSprop optimization algorithm^42,43^ with a learning rate of 0.001 and minibatch size of 64 for 100 epochs (cycles of DL training) until convergence of the loss function and variance-bias tradeoff (Figure S1).^22^ CVAE models were implemented using the mdlearn Python library.^15^ Full details of the training datasets are as follows.

#### Pre-trained DL WE simulations

For these simulations, a deep CVAE model was pretrained on representative conformations of each MSM microbin for NTL9 protein folding with the addition of 21 folded-state conformations. These conformations were generated using 21 1-ns of cMD simulations propagated from the single folded structure of our MSM. Given that the MSM only included a single folded conformation, the addition of 21 folded conformations was necessary to provide an equal number of conformations for the folded and unfolded states in the training set for the CVAE model (i.e., 22 conformations for each state).

#### On-the-fly DL WE simulations

For these simulations, an initial CVAE model was trained on a base dataset of 2000 conformations from 20 ns (2000 steps) of synMD trajectories combined with the 22 folded conformations mentioned above. A new CVAE model was then trained every 10 WE iterations by updating the base dataset with data (contact maps) from the latter 50 WE iterations. This periodic updating of the training dataset enabled the CVAE model to “learn” an improved internal latent space representation of the system as new regions of conformational space were explored.

### Binless control simulations with sorting by RMSD

To determine the impact of DL on the efficiency of our WE simulations, we carried out our binless control WE simulations without the use of DL (CVAE model) to identify outlier trajectories. The trajectories in these simulations were first randomly shuffled before applying the WE resampling procedure based on the C_α_ RMSD from the folded structure. We also assessed the effectiveness of sorting the trajectories solely by the LOF score in CVAE latent space representations (with no additional ranking by C_α_ RMSD from the folded structure).

### Binned control simulations with RMSD progress coordinate

As another point of comparison, we ran binned control WE simulations without the use of DL, employing a onedimensional progress coordinate consisting of the C_α_ RMSD from the folded structure and rectilinear bins positioned using the minimal adaptive binning (MAB) scheme. ^44^ We applied the MAB scheme with 10 rectilinear bins between the trailing and leading trajectories, up to 2 bins for the bottleneck and leading trajectories, and 6 target trajectories per bin to yield a similar total number of trajectories as the other WE protocols used in this study (*M* = 72 trajectories).

### Calculation of the folding rate constant

The folding rate constant *k_fold_* was directly calculated from our WE simulations using the following exact Hill relation:^45^

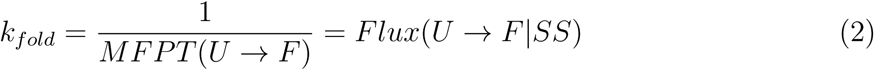

where *MFPT* (*U → F*) is the mean first-passage time (average time) it takes for the protein to transition from the unfolded to the folded state and *Flux*(*U → F|SS*) is the nonequilibrium steady state probability flux carried by trajectories originating from the unfolded state and reaching the target folded state Uncertainties represent 95% credibility regions over 10 trials of WE simulation, as determined using a Bayesian bootstrap method.^46,47^ The ground-truth *k_fold_* value was determined from our generative MSM model using the Deeptime Python library.^48^

### Estimating DL-enhancement of WE efficiency

The efficiency *S_k_*of a DL-enhanced WE simulation over a control WE simulation in computing a rate constant of interest (here, the folding rate constant *k_fold_*) was estimated using the following equation:^16,49^

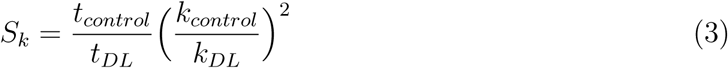

where *t_control/test_* is the total simulation time for a control/test simulation, respectively, and *k_control/test_* is the relative error in the *k_fold_*estimate (ratio of the width of the uncertainty of the rate constant relative to the value of the rate constant, where the uncertainty represents the 95% credibility region) for the corresponding simulations. Thus, the efficiency of a WE simulation in calculating the rate constant is determined by taking the ratio of the total simulation times for the control and test WE protocols that would be required to estimate the rate constant with the same relative error, assuming that the square of the width of the 95% credibility region on the rate constant is inversely proportional to the total simulation time.^49^

## 3 Results and Discussion

We have developed a WE simulation method that applies DL to learn an effective progress coordinate “on-the-fly” during a simulation. The DL process involves identifying outlier trajectories based on a Local Outlier Factor (LOF) anomaly score in latent space and the C_α_ RMSD from the target state in real space. Our benchmark application is the simulation of the millisecond folding process of the NTL9 protein using discrete-state synMD trajectories. To assess the impact of DL on the efficiency of the WE simulations, we ran control WE simulations without DL using (i) a “binless” approach where trajectories are sorted by the C_α_ RMSD from the folded state, and (ii) a rectilinear, adaptive binning approach along a one-dimensional progress coordinate consisting of the C_α_ RMSD from the reference folded structure. We also determined the effectiveness of applying DL on-the-fly during a WE simulation vs. pre-training on cMD simulation data prior to running a WE simulation. Key details of all WE simulation protocols used in this study are summarized in Table 1.

**Table 1:** WE simulation protocols used in this study. For each WE protocol, we summarize the criteria for identifying outlier trajectories and simulation data used for DL training. WE simulations using either pre-trained or on-the-fly DL identified outlier trajectories in a “binless” manner based on the LOF score in a three-dimensional CVAE latent space model of the system and C_α_ RMSD from the folded structure in real space. Two types of control simulations were run without the use of DL: (i) binless control simulations where outlier trajectories were identified based on the C_α_ RMSD from the folded structure, and (ii) binned control simulations where adaptive binning was applied along a progress coordinate consisting of the C_α_ RMSD from the folded structure.

| WE protocol | Outlier identification | Deep-learning (DL) training |
| --- | --- | --- |
| On-the-fly DL | By LOF and RMSD | Every 10 WE iterations on data from the latter 50 WE iterations |
| Pre-trained DL | By LOF and RMSD | Once from 2.5 μs cMD simulations |
| Binless Control | By RMSD | None |
| Binned Control | By RMSD | None |

### Unsupervised learning identifies unfolded, intermediate, and folded states

Before applying on-the-fly DL during a WE simulation, we verified that a CVAE latent space representation of data from a set of cMD simulations of the NTL9 folding process (2.5 µs of total simulation time) could identify key stable or metastable states. As shown in Figure 3, a three-dimensional CVAE representation of the simulation data was sufficient for this identification when the data was colored according to the C_α_ RMSD from the folded structure.

**Figure 3:**
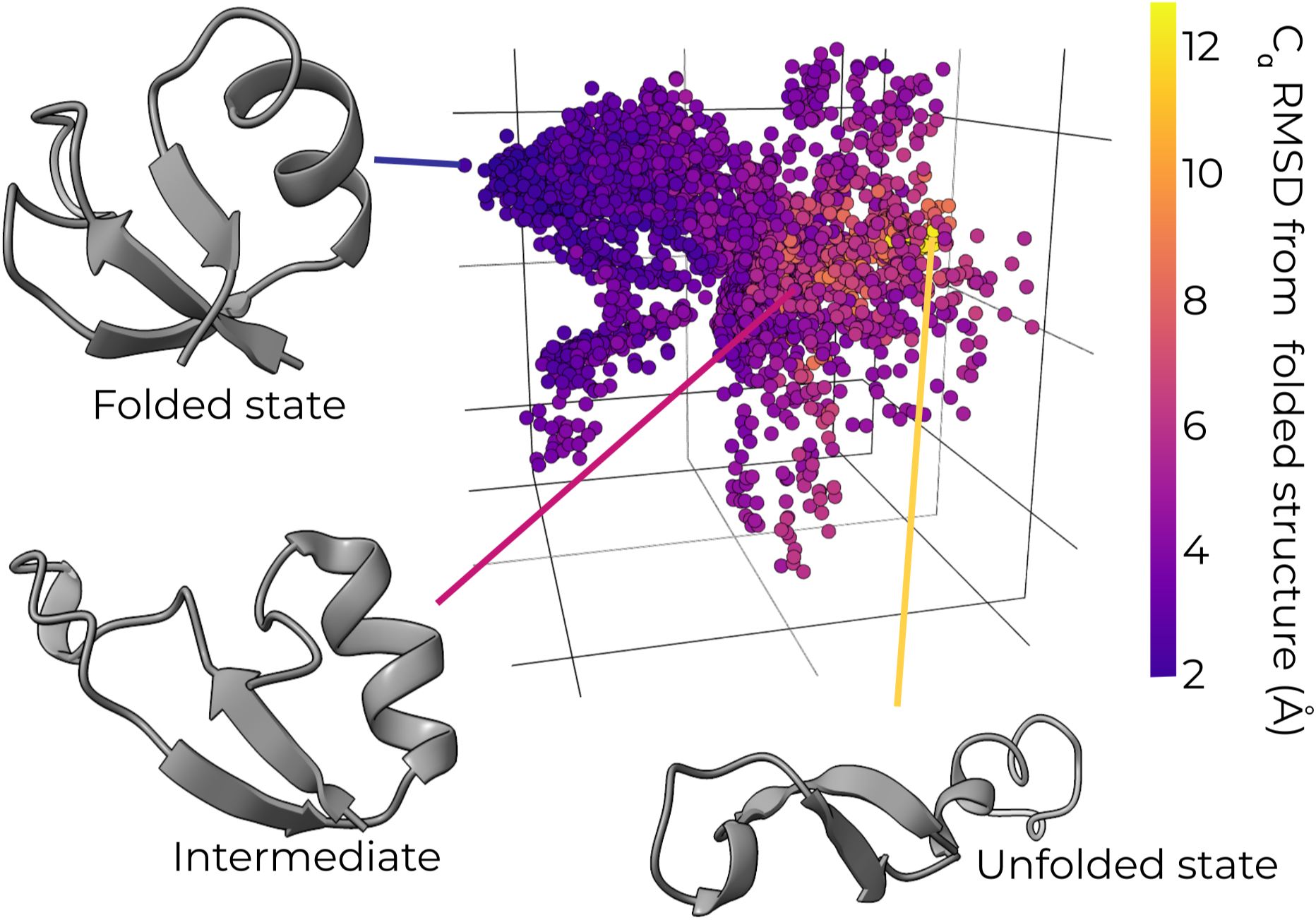
Pre-trained CVAE model identifies key states for NTL9 protein folding process. A three-dimensional CVAE latent space model pre-trained using a NTL9-folding simulation dataset with data points colored by the C_α_ RMSD from the folded structure. The training dataset was generated using a set of representative structures for the microbins of a MSM (one structure for each microbin) that was constructed using 2.5 µs total simulation time of cMD simulations with conformations saved every 10 ps and an additional 21 folded state structures generated from 21 ns of cMD simulations from a folded state structure. This pre-trained CVAE model separates key states of the NTL9 folding process, revealing unfolded, intermediate, and folded states.

### A real-space structural metric is necessary to identify outliers

Our results revealed that the sorting of trajectories by LOF score in latent space was not sufficient for efficient generation of successful folding events and that additional sorting using a real-space structural metric (i.e., RMSD) was necessary. When only sorting by the LOF score, the WE simulations sampled primarily the unfolded state (high-RMSD region) (Figure 4A). On the other hand, additional sorting by RMSD resulted in extensive sampling of latent space and the identification of outlier conformations along the periphery (Figure 4B). This additional sorting more than doubles the number of successful folding events by replicating trajectories at the leading edge while terminating trajectories at the trailing edge (Figures 5A and S2). Furthermore, binless control simulations (without DL) with sorting of trajectories by only RMSD, were able to generate successful events while those with random sorting of trajectories were unable to generate any successful events (Figures S3-S4).

**Figure 4:**
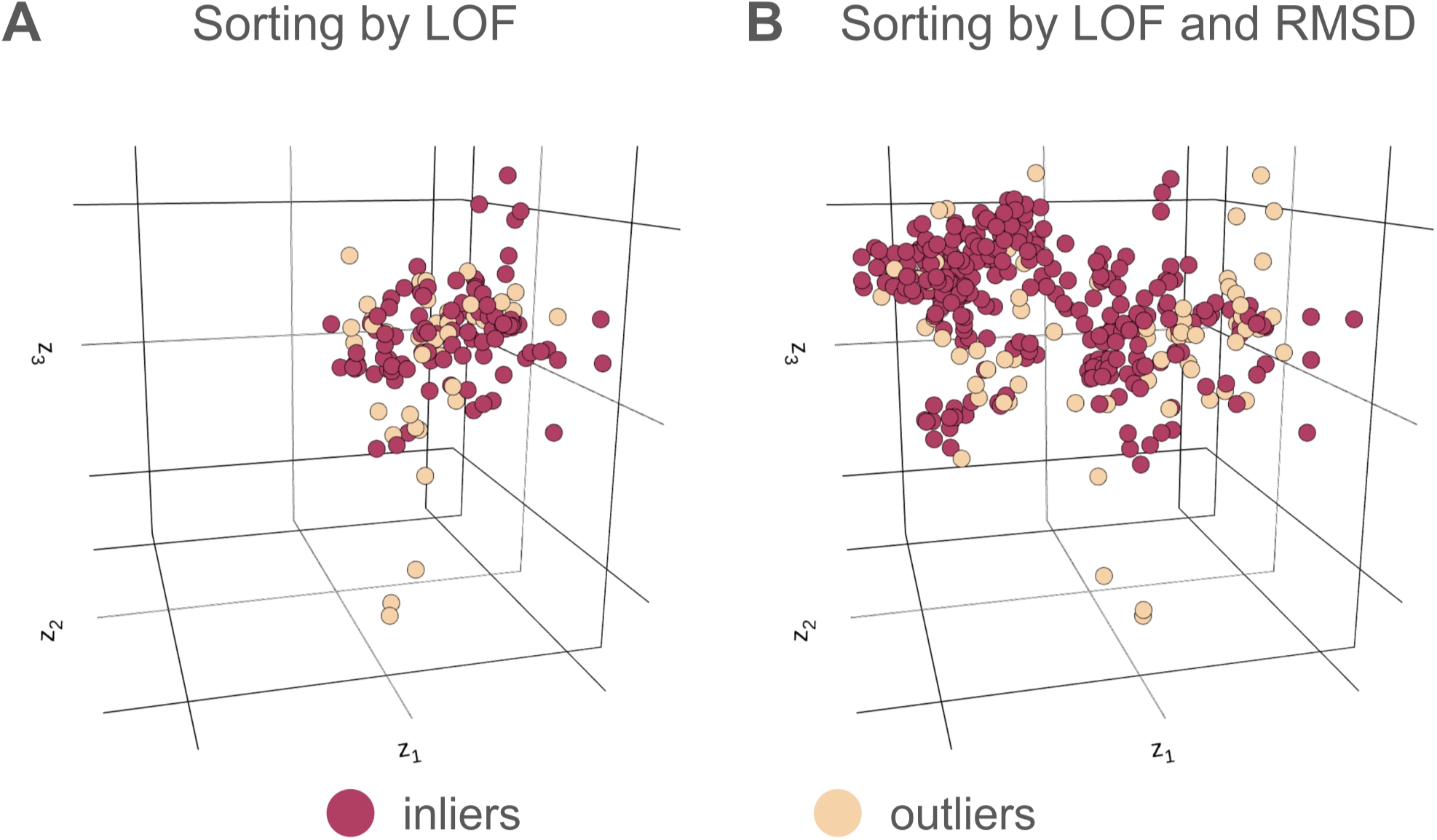
A real-space structural metric is necessary to identify productive outliers in latent space. Three-dimensional CVAE latent space representations of the NTL9 folding process based on pre-trained DL with A) sorting by only LOF score and B) sorting by both LOF score and a real-space metric (*C*_α_ RMSD from the folded structure). Conformations identified as outliers are colored yellow and those identified as inliers are colored red.

**Figure 5:**
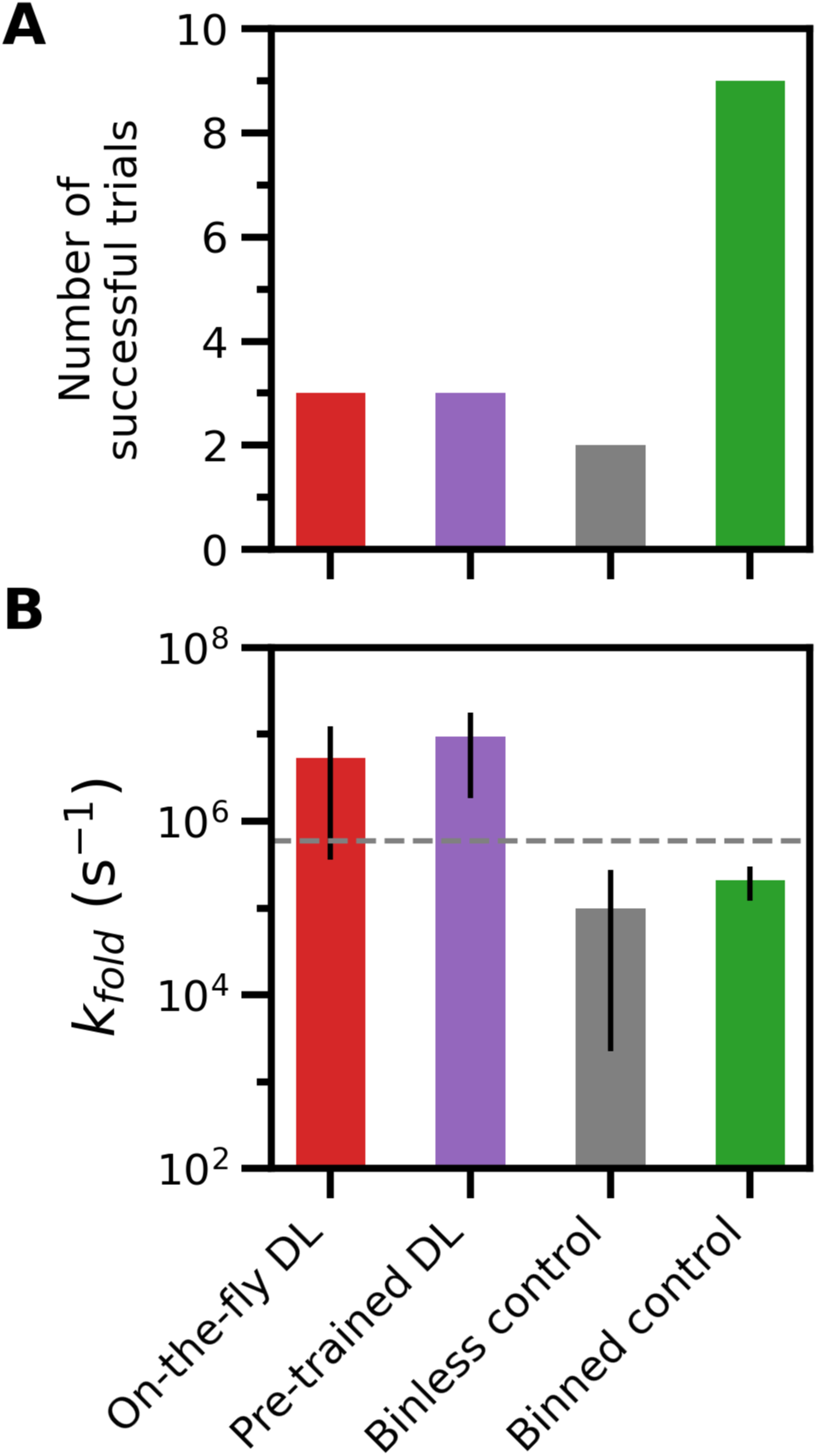
Number of successful simulation trials and average. *k_fold_***estimates generated by each WE protocol.** (A) Number of successful trials for each WE protocol. A trial was considered successful if the *k_fold_* estimate was within one order of magnitude of the ground-truth value (dashed horizontal line in (B)). (B) The average *k_fold_* estimate generated by each WE protocol. Uncertainties represent 95% credibility regions over 10 trials for each WE protocol, as determined using a Bayesian bootstrap method.^46,47^ Data shown for each WE protocol is based on the same total simulation time of 14.5 µs.

### On-the-fly DL enhances WE efficiency

We next tested the effectiveness of onthe-fly DL of a progress coordinate during a WE simulation of the NTL9 folding process. Compared to the binless control simulations, on-the-fly DL simulations were >3-fold more efficient in estimating a kfold value (Table 2). Pre-trained DL simulations were also more efficient, but to a smaller extent (1.7-fold), partially due to the fact that our estimate of the efficiency includes the 2.5 µs in aggregate of cMD simulations used for DL training. Both on-the-fly and pre-trained DL simulations exhibited a substantially lower variance in the rate-estimates relative to the binless control simulations with the same total simulation time (Figures 5B and S5).

**Table 2:**
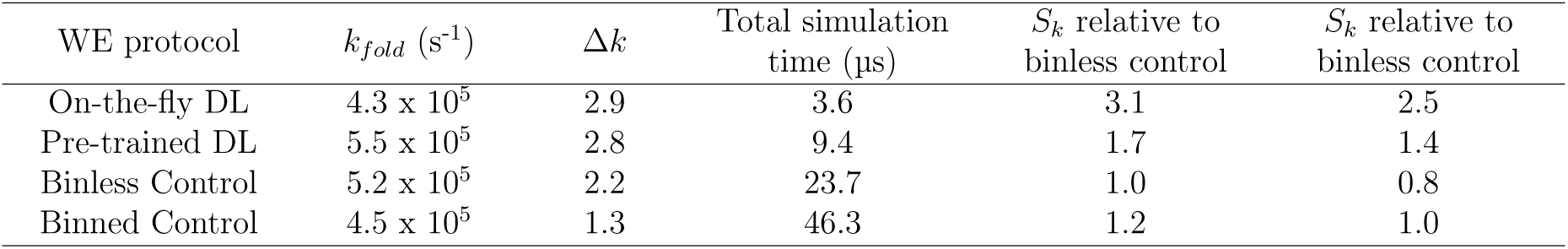
Efficiency of DL-enhanced vs. control WE simulations. The efficiency *S_k_*is estimated by taking the ratio of total simulation times for the DL-enhanced vs. binless or binned control WE simulations that would be required to estimate the rate constant kfold with the same relative error Δ*_k_*, which is the ratio of the width of the 95% credibility region on kfold and the estimated value of *k_fold_*(see Methods).^16,49^ The total simulation time for pre-trained DL simulations includes time invested for the cMD simulations used for DL training. All simulations were run until the ground-truth value fell within their corresponding 95% credibility regions (Figure S7).

| WE protocol | $k_{fold}$ ($s^{-1}$) | $\Delta k$ | Total simulation time ($\mu s$) | $S_k$ relative to binless control | $S_k$ relative to binned control |
| --- | --- | --- | --- | --- | --- |
| On-the-fly DL | $4.3 \times 10^5$ | 2.9 | 3.6 | 3.1 | 2.5 |
| Pre-trained DL | $5.5 \times 10^5$ | 2.8 | 9.4 | 1.7 | 1.4 |
| Binless Control | $5.2 \times 10^5$ | 2.2 | 23.7 | 1.0 | 0.8 |
| Binned Control | $4.5 \times 10^5$ | 1.3 | 46.3 | 1.2 | 1.0 |

We also compared the efficiency of our DL-enhanced WE simulations relative to binned control WE simulations employing the MAB scheme (see Methods),^44^ which has been shown to efficiently surmount large barriers. We applied this adaptive binning scheme along a one-dimensional progress coordinate consisting of the C_α_ RMSD from a reference folded structure. The use of DL also showed a marginal increase in efficiency compared to the binned control simulations, with a 2.5-fold gain for on-the-fly DL and 1.4-fold gain for pretrained DL (Table 2). Among all the WE protocols, the adaptively binned WE simulations were the most efficient in generating initial folding events (Figure S2), but did not reach a steady-state that yields the ground-truth *k_fold_* value. The DL-enhanced WE simulations were reasonably converged, reaching the ground-truth value within the same total simulation time.

Compared to the binned control simulations, the greater efficiency of both the on-the-fly and pre-trained DL-enhanced WE simulations in reaching the ground truth appears to be due to their “binless” nature. These binless strategies allow us to allocate a majority of the M trajectories for exploitation towards the target state, potentially leading to faster convergence to a steady state that yields the ground-truth *k_fold_*value. However, these strategies resulted in a relatively wide range of trajectory weights, yielding abrupt “rampup” times in the kinetics in contrast to the exponential ramp-up times that are characteristic of a binned strategy (Figures S5-S6),^50^ and thereby relatively large variances in the folding rate estimates between trials (Figure 5A). On the other hand, the WE simulations with adaptive binning resulted in a more narrow range of trajectory weights (Figure S8) with slower convergence to a steady state, but lower variance in rate estimates between trials.

### Overhead of DL training

We note that the reported efficiencies (*S_k_* values) for our DL-enhanced WE simulations do not include the overhead for training the CVAE model.

With the exception of the pre-trained DL protocol, a single trial of each WE protocol was completed within minutes to hours, highlighting the advantage of using synMD trajectories for rapid testing in methods development. Although the wall-clock time for a pre-trained DL simulation was only 1.15 hrs for running the weighted ensembles of synMD trajectories, *∼*30 hrs was required to complete the cMD simulations for pre-training the CVAE model (Table 3). On the other hand, the on-the-fly DL simulations required only a small initial dataset (here, 20 ns of synMD trajectories). Relative to the binless and binned control simulations, the >35-fold longer wall-clock time of the on-the-fly DL simulations is due to the substantial overhead for training the deep CVAE models. For future simulation studies, we recommend starting with on-the-fly DL WE simulations to generate initial successful pathways for a rare-event process of interest, then running additional WE trials using the final updated DL model.

**Table 3:** Wall-clock times for each WE protocol. Wall-clock times required for running a single WE trial simulation with 1.45 µs total simulation time and any DL training. Each simulation was run using a single thread of an AMD Ryzen 9 7950X CPU. DL training was performed on a single NVIDIA RTX 4090 GPU.

| WE protocol | Wall-clock time (hrs) |
| --- | --- |
| On-the-fly DL | 2.85 |
| Pre-trained DL | 31.15 |
| Binless Control | 0.08 |
| Binned Control | 0.05 |

## 4 Conclusions

We have developed a weighted ensemble (WE) path sampling method that applies deep learning (DL) on-the-fly to learn effective progress coordinates during a simulation. Our DL enhanced WE method learns progress coordinates by identifying outlier trajectories based on relatively low local densities in latent space, as quantified by local outlier factor (LOF) scores, and structural information in real space (RMSD from the target structure). We applied our method to simulations of the millisecond NTL9 protein folding process using discrete-state synthetic molecular dynamics trajectories.

Our “binless” WE method was >3-fold more efficient than binless control simulations with no DL and 2.5-fold more efficient than binned control simulations with no DL. These gains in efficiency underscore the value of projecting high-dimensional simulation data onto a low-dimensional latent space model for identifying progress coordinates that are effective for rare-event sampling. It is worth noting that our reported efficiency gains account for only the total simulation times and not the overhead of training the DL models. To reduce this overhead, we have been integrating the WESTPA software with the Colmena framework^51,52^ to implement model-training in parallel with the execution of WE simulations (unpublished work).

While our binned control simulations achieve the highest precision in rate-constant estimates, these simulations do not reach the ground-truth rate constant within the same total simulation time as that used for our on-the-fly DL protocol. On the other hand, the on-the-fly DL protocol reaches the ground truth, but with a higher variance in the rate-constant estimates. Finally, the necessity of using a real-space RMSD metric in addition to the latent space LOF score highlights the challenge of identifying productive outlier conformations in latent space without a physically intuitive structural metric. In future work, we will develop generative models, i.e., based on emerging neural operators^53^ to identify outlier trajectories for replication. It is our hope that these models will also reduce the variance among individual trials of WE simulation.

## Acknowledgement

We thank John Russo and Daniel Zuckerman (OHSU) for making available their synMD model of the NTL9 folding process. JMGL was supported by a Molecular Sciences Software Institute Predoctoral Fellowship under NSF grant CHE-2136142. AB and AR were supported by National Institutes of Health Award Number P01AI165077 and the Coalition for Epidemic Preparedness Innovations (CEPI). Funding was also provided to LTC by NIH grant R01 GM1151805 and a “Characteristic Science Applications” subaward from the Texas Advanced Computing Center (TACC) under NSF grant 2139536. We are grateful for assistance from TACC consultants, Kent Milfield and Albert Lu.

## Supporting Information Available

Supplementary figures and tables. All input files and scripts needed to run and analyze the WE simulations in this study are provided in the GitHub repository: https://github.com/ westpa/DL-enhancedWE and deposited on Zenodo under DOI: https://doi.org/10.5281/ zenodo.13387514.

The following files are available free of charge.

SI: Supplementary figures

## TOC Graphic

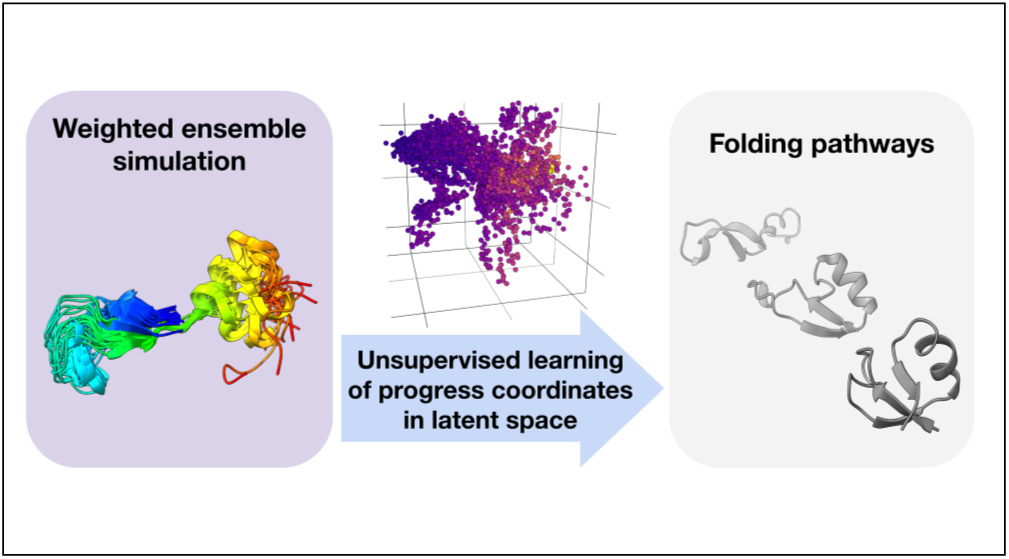

