## Supporting Information for "Unsupervised learning of progress coordinates during weighted ensemble simulations: Application to millisecond protein folding"

### 1 Supplementary FIgures

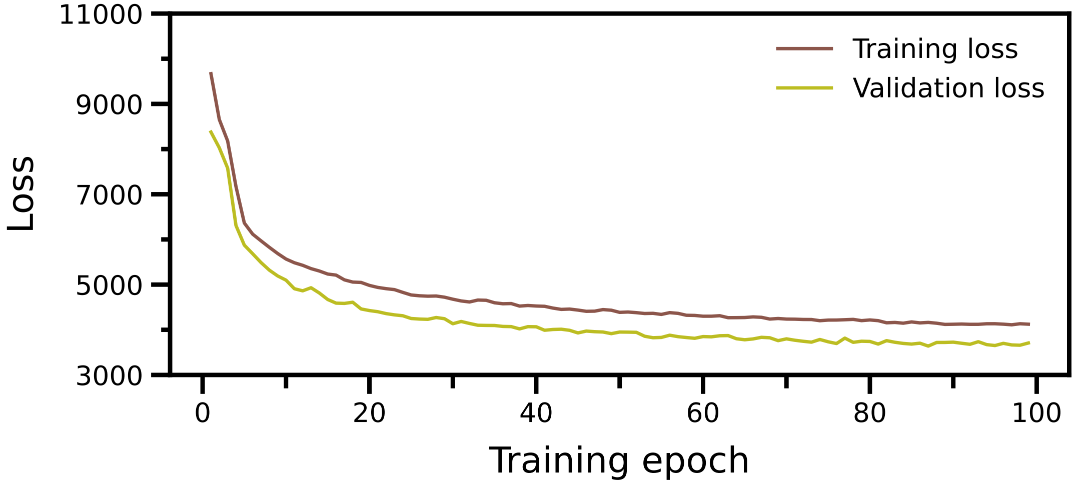

Figure S1: **Training loss as a function of training epoch for the pre-trained CVAE model.** Training and validation losses for the CVAE model converged after 100 training epochs (cycles of DL training). The training loss appears higher than the validation loss because the latter was measured at the end of an epoch whereas the former was averaged over each training step of the epoch.

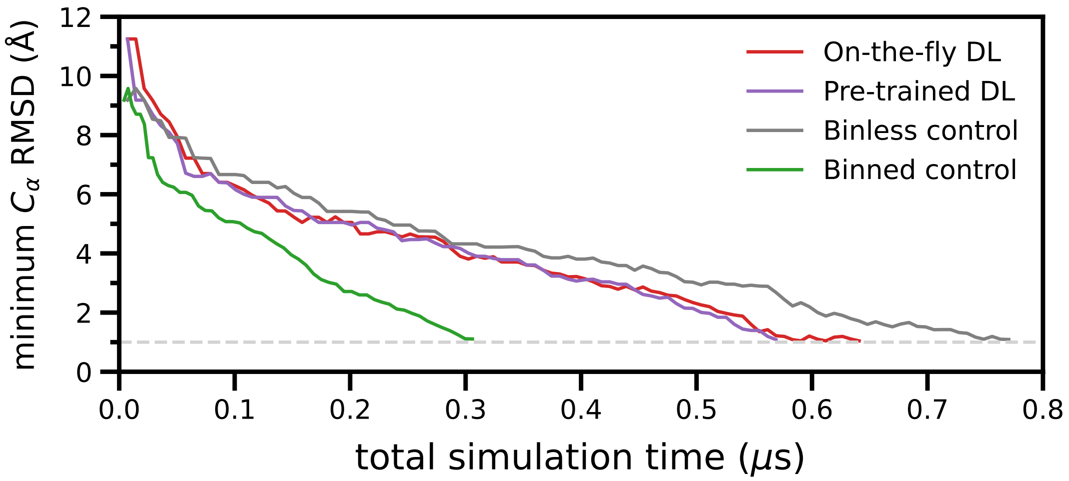

Figure S2: **Time-evolution of the minimum $C_\alpha$ RMSD from the folding structure reached by each WE protocol.** The data trace for each WE protocol was truncated at the total simulation time where a folding event was first generated (dotted line).

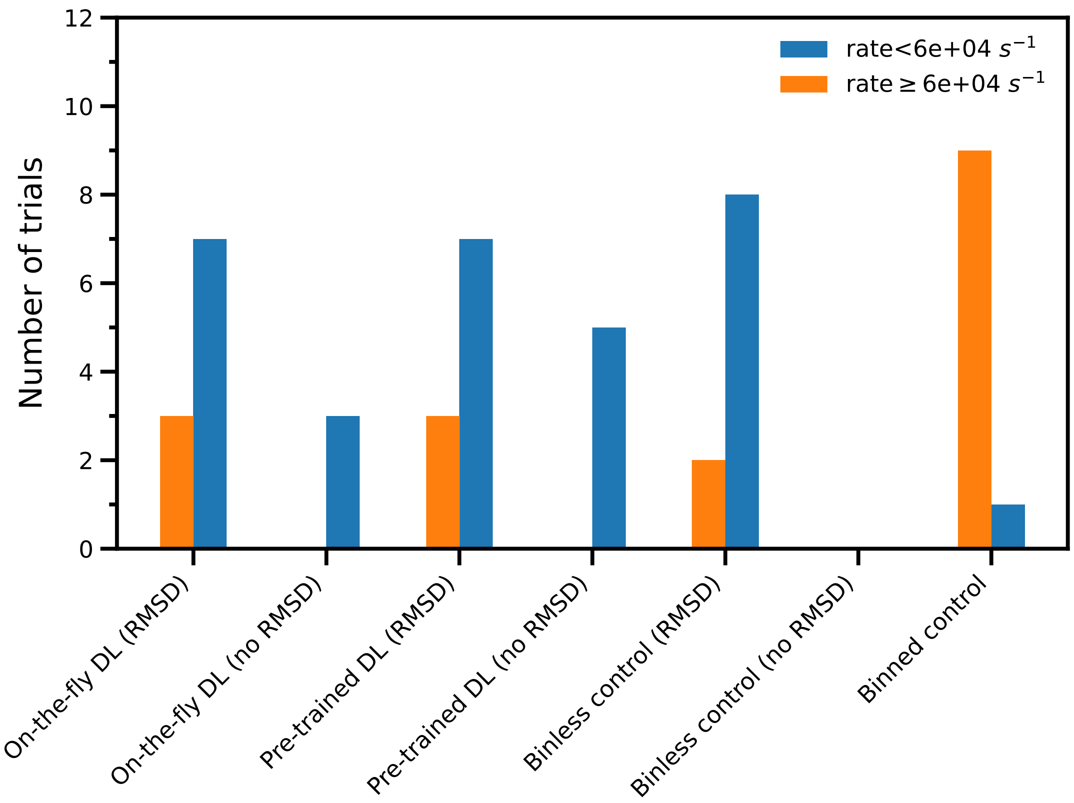

Figure S3: **Number of successful trials for each WE protocol.** Orange bars indicate trials with folding-rate constant $k_{fold}$ estimates within one order of magnitude of the ground-truth value, whereas blue bars indicate trials that have not yet reached the ground-truth value (beyond one order of magnitude). The use of a real-space structural metric (i.e., $C_\alpha$ RMSD from the folded structure) to further sort trajectories prior to WE resampling yielded a greater number of trials with successful events.

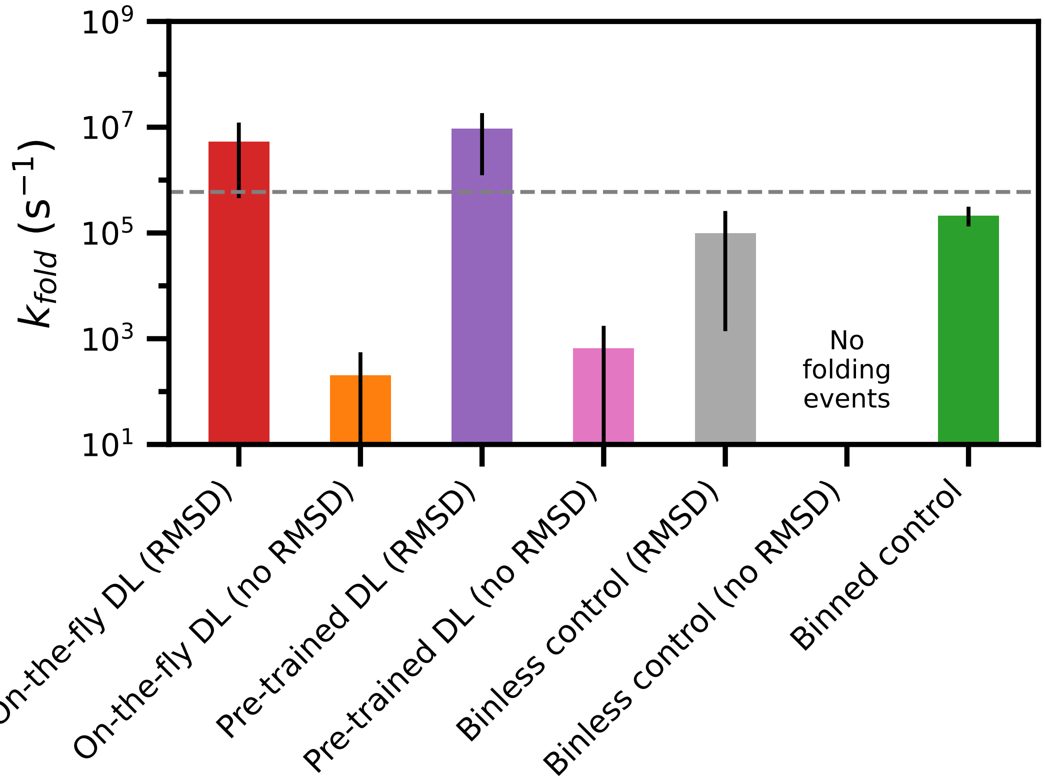

Figure S4: **Average folding-rate constant $k_{fold}$ estimates for each WE protocol.** Uncertainties represent 95% credibility regions over 10 trials with each WE protocol, as determined using a Bayesian bootstrap method.[1,2] The ground-truth value is shown as the dashed horizontal line. The total simulation time for each simulation protocol was 14.5 µs.

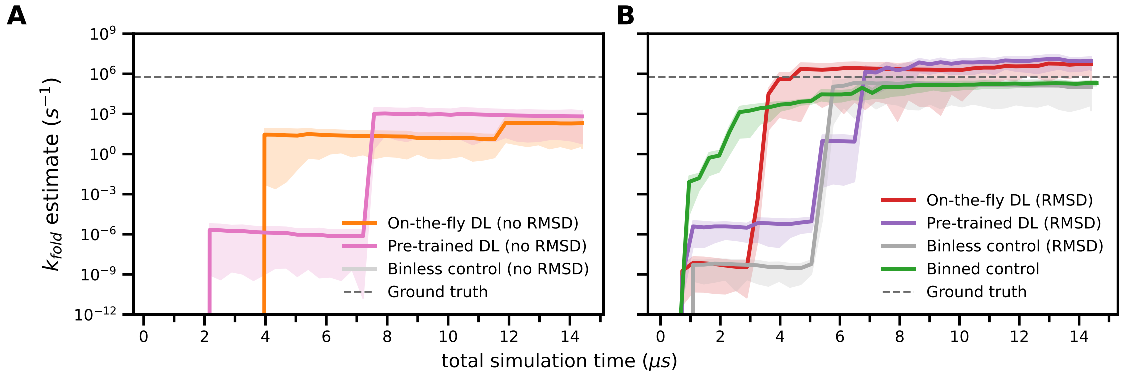

Figure S5: **Time-evolution of the average folding-rate constant $k_{fold}$ estimate for each WE simulation protocol.** (A) Folding-rate constant $k_{fold}$ estimate of three protocols with random sorting of trajectories (instead of sorting by real-space RMSD) prior to WE resampling. Binless control simulations (without RMSD sorting) were unable to generate any successful folding events within a total simulation time of 14.5 µs. (B) Folding-rate constant $k_{fold}$ estimates using four WE protocols with sorting of trajectories by real-space RMSD prior to WE resampling.

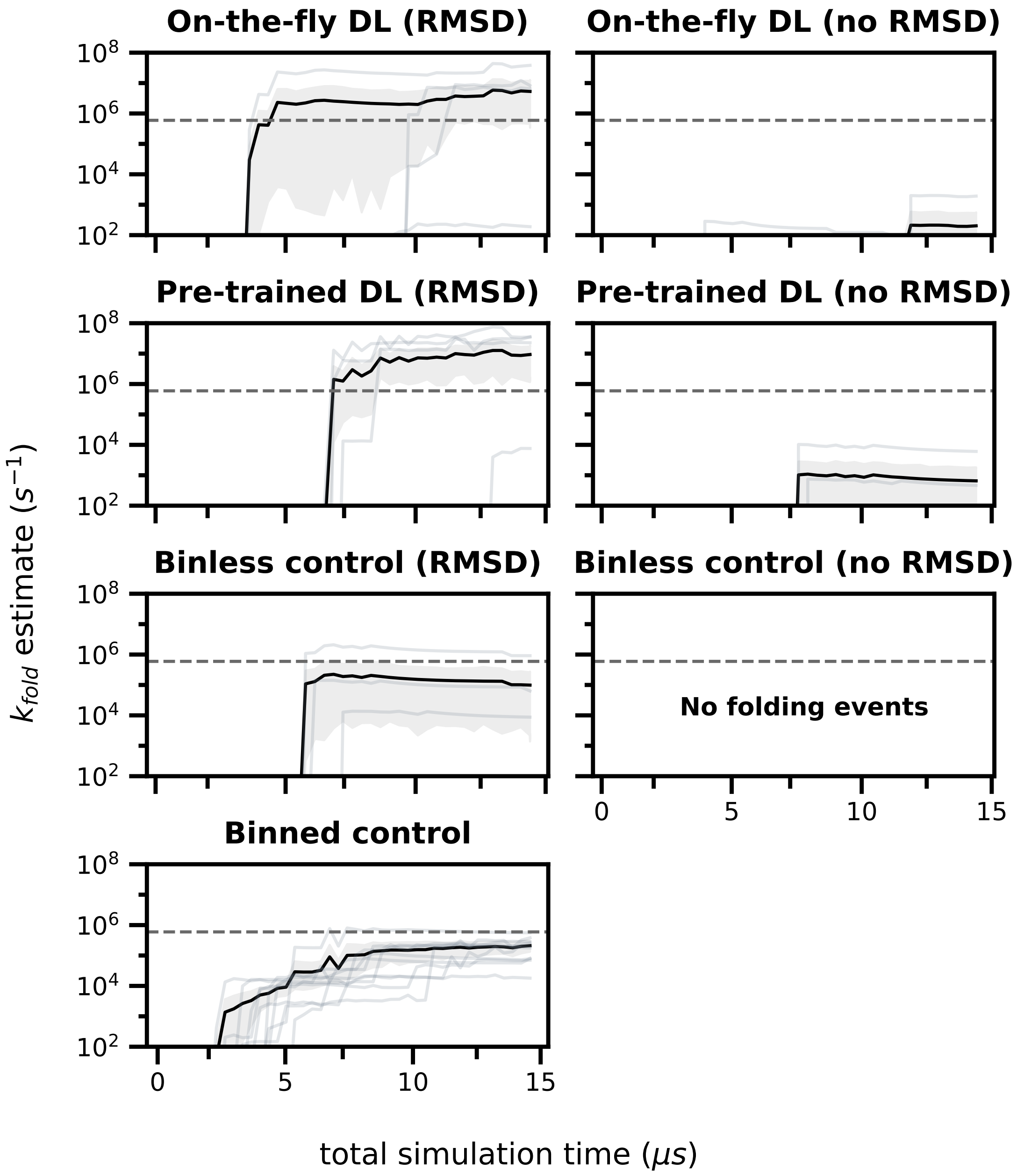

Figure S6: **Time-evolution of the folding-rate constant $k_{fold}$ estimate for each of the 10 simulations run for each WE protocol.** Each panel represents the folding-rate constant estimation for a single WE protocol. The gray traces represent individual trials and the black trace represents the average of the 10 trials with a 95% credibility region (gray shading). The horizontal dashed line represents the ground-truth value.

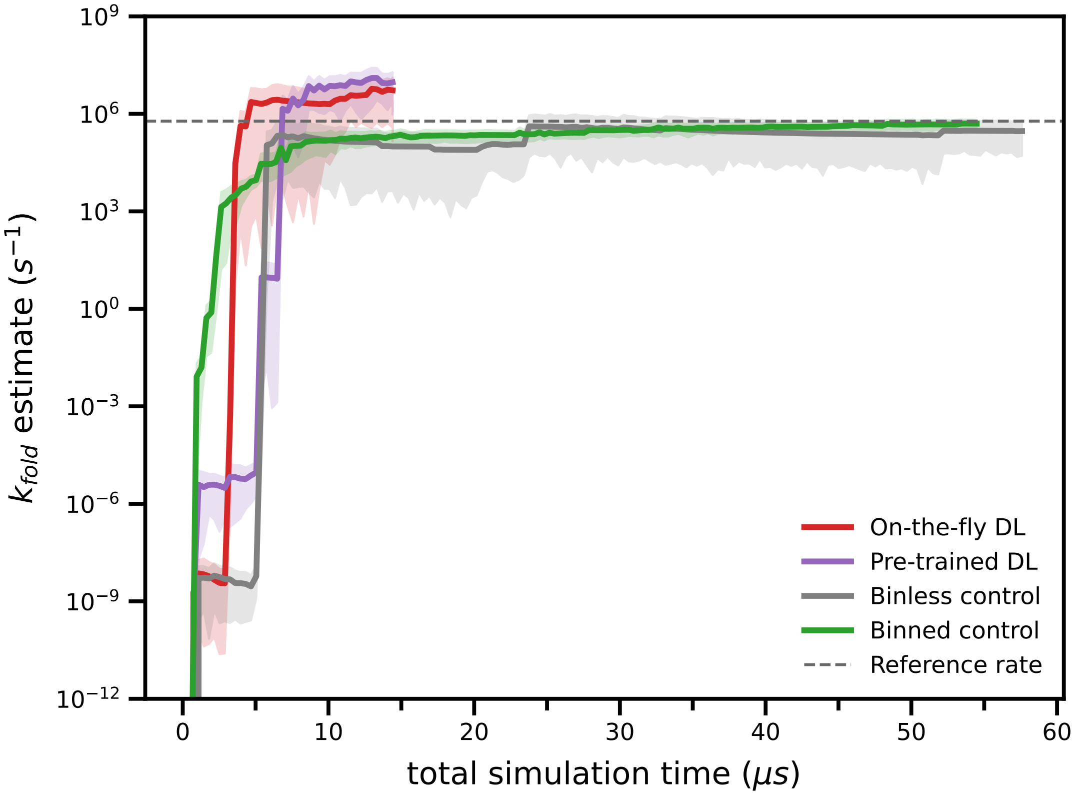

Figure S7: **Time-evolution of the average folding rate-constant $k_{fold}$ estimate.** All simulations were extended until reaching the ground-truth value for the folding-rate constant $k_{fold}$. Compared to the DL methods, binned control simulations required 5-12x the amount of total simulation time and reached the ground-truth value with a lower uncertainty to reach the ground-truth value (95% credibility region).

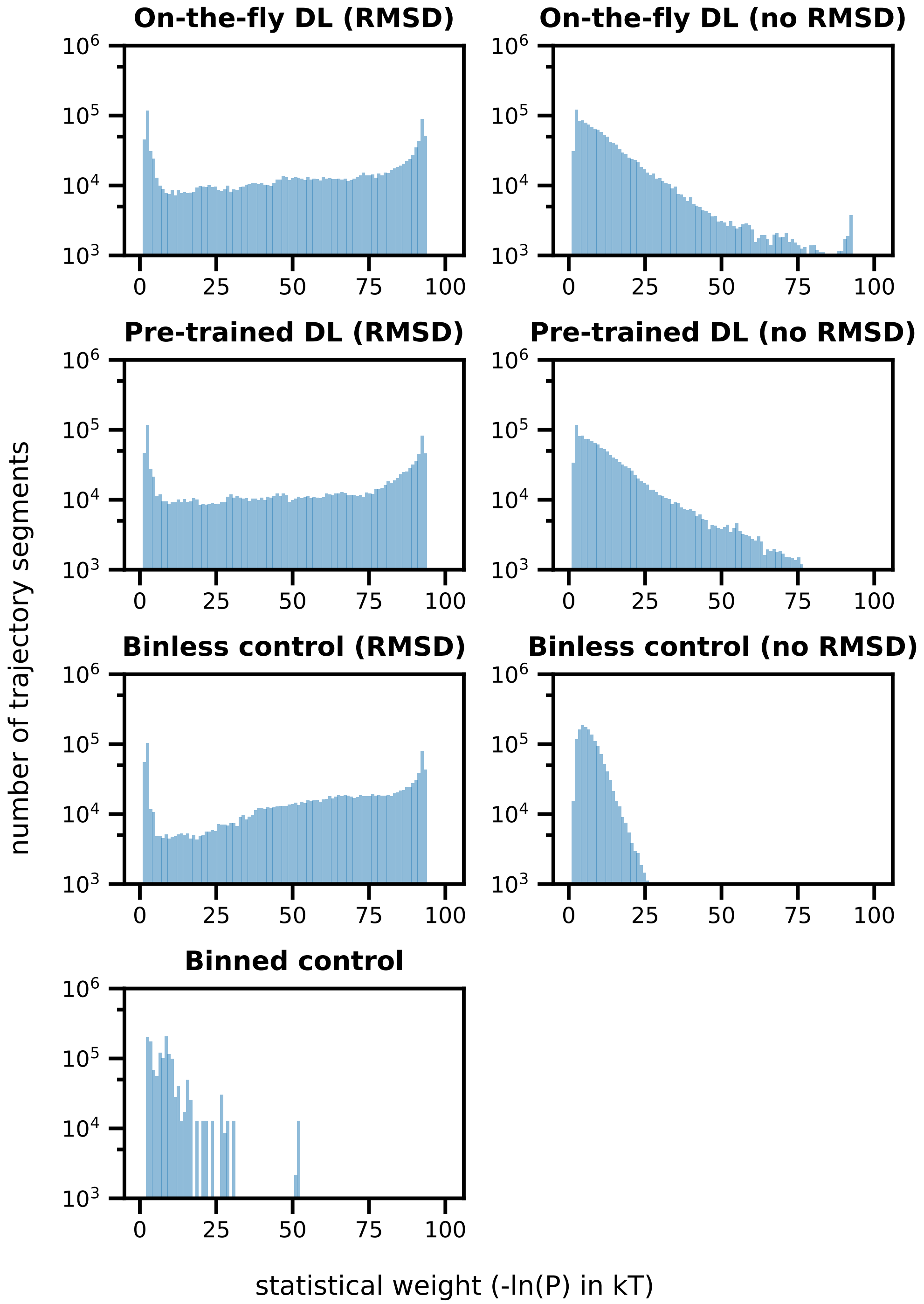

Figure S8: **Histogram of trajectory weights for each WE protocol.** With the exception of the binless control (no RMSD) simulations, the "binless" resampling protocols (top three rows) generated a wide range of trajectory weights. In contrast, the binned control simulations generated a narrower range of trajectory weights.
